# Shared brain transcriptome signatures of social evolution and plasticity in shell-dwelling cichlid fishes

**DOI:** 10.64898/2026.11.07.765067

**Authors:** Pol Sorigue, Alex Jordan, Walter Salzburger, Rui F. Oliveira

**Author notes:** Email: Pol Sorigue; Alex Jordan; Walter Salzburger; Rui Oliveira.

## Abstract

Inter-individual variability in phenotypic traits is widespread in nature both across and within species, with similar patterns of variation often occurring between and within species. Given that these two types of variation are generated by evolutionary and plasticity processes respectively, it raises the question to what extent the causal mechanisms underlying such variability at these two different levels are shared or not. To address this question, we studied the variation in brain transcriptomes in Tanganyikan cichlids that exhibit both intra- and interspecific variation in social behavior. We compared two facultatively social cichlid species, each containing naturally occurring solitary and social-living individuals, with a closely related obligatory solitary resembling the ancestral state. Differential gene expression and Gene Set Enrichment Analyses were performed in the telencephalon and diencephalon transcriptomes to investigate the molecular basis of both behavioral plasticity and the evolution of sociality. Behavioral plasticity involved few differentially expressed genes but extensive changes in biological pathways, supporting a highly polygenic architecture of social behavior. Across both species, social-living individuals consistently upregulated pathways involved in translation, protein homeostasis, and energy metabolism, whereas solitary individuals showed enrichment of neuronal morphogenesis, cell adhesion, and structural plasticity. Remarkably, these same functional programs were associated with the independent evolution of sociality, revealing a strong overlap between evolutionary and plastic transcriptional responses. This convergence was largely restricted to the telencephalon, while the diencephalon showed limited and inconsistent functional signatures. Together, our results support the repeated recruitment of gene regulatory programs during the evolution of sociality and demonstrate that behavioral plasticity and evolutionary transitions share common neurogenomic responses, consistent with a conserved genetic toolkit for social behavior.

## Introduction

Becoming social is one of the major transitions in the history of life on Earth (Szathmáry & Smith, 1995). By associating with conspecifics (‘social-living’), social individuals gain fitness advantages such as reduced predation risk, improved foraging, and access to social information, while also facing costs, including resource competition, increased exposure to pathogens, and social stress (Krause & Ruxton, 2002). Social-living has evolved multiple times in animals and across a wide range of environments, and animal societies vary greatly in size and complexity (M. Taborsky et al., 2021). Unlike in solitary organisms, individuals in social groups must continuously adjust their behavior in response to other individuals of their own species, making survival and reproductive success at least partly dependent on the actions and states of conspecifics. This interdependence on conspecifics comes with particular challenges, as fitness is no longer determined solely by interactions with the physical environment and species interactions, but also by the complexity of the intraspecific social environment (B. Taborsky & Oliveira, 2012). Navigating this reliance on others introduces an additional layer of behavioral and cognitive complexity, forcing individuals to cope with shifting social roles, maintain hierarchies, and integrate multi-sensory social information in real time, in addition to all other ecological needs shared with non-social individuals (e.g. feeding, avoiding predators, finding mates)

Behavior has been biologically grounded to the brain and therefore variation in behavior across social contexts has been linked to changes in the brain at multiple levels (Fernald, 2015; Robinson et al., 2008). At the structural level the challenges imposed by social living are thought to have driven the evolution of bigger and more complex brains, as more sophisticated neural mechanisms are required to process, predict, and respond to social information (Dunbar, 1998; Reader & Laland, 2002). In particular, an evolutionary conserved ‘social decision-making network’ has been identified across vertebrates, consisting of a set of interconnected brain regions that integrates social information, internal physiological state, and environmental cues to regulate social behavior (Goodson, 2005; O’Connell & Hofmann, 2012). At the functional level, constitutive differences in brain transcriptomes and epigenomes directly correlate with distinct social phenotypes (e.g. Cardoso et al., 2018; Gospocic et al., 2021; Jones et al., 2017; Simola et al., 2016; Young et al., 2019), and shifts in neurogenomic profiles parallel the behavioral flexibility needed to navigate changing social environments (e.g. Gospocic et al., 2017; Renn et al., 2008; Rittschof et al., 2014; Saul et al., 2017; Simões et al., 2015). Consequently, social behavior induced changes in gene expression provide a flexible molecular substrate through which individuals can modify neural function and behavior in response to dynamic social environments (Cardoso et al., 2018; Oliveira et al., 2016). Together, these findings indicate that specific brain gene expression profiles in the brain parallel specific behavioral states and that context-dependent changes in behavioral states (i.e. behavioral plasticity) are accompanied by shifts between brain transcriptomic profiles (Cardoso et al., 2015) Thus, brain transcriptomes are a valuable readout for behavioral variation and provides a functional link between neurogenomic and behavioral states. Variation in gene expression also represents a potential molecular link between genetic variation and phenotypic plasticity. Gene expression levels can themselves be heritable phenotypic traits, with genetically determined differences in expression contributing to phenotypic variation among individuals (Hill et al., 2021; Zheng et al., 2011). At the same time, gene expression is highly responsive to environmental conditions and can therefore constitute a key component of phenotypic plasticity (Chevin et al., 2022). Thus, variation in gene expression may arise both from genetic differences among individuals and from environmentally induced changes within individuals, suggesting that the molecular mechanisms underlying heritable phenotypic variation and phenotypic plasticity in social behavior may be at least partially shared.

Social phenotypes typically emerge from the coordinated regulation of gene sets rather than from the action of single ’social genes’ (Kapheim et al., 2015; Renn et al., 2016). Research across diverse taxa suggests that the evolution of sociality often involves the co-option of conserved genes and gene regulatory networks that are repeatedly recruited to generate similar phenotypes across independent evolutionary lineages (Rittschof et al., 2014; Rittschof & Robinson, 2016; Schumer et al., 2011; Toth & Robinson, 2007). For example, distantly related vertebrates that have independently evolved social monogamy exhibit conserved neural transcriptomic signatures with shared gene-ontology categories, and/or even homologous genes suggesting a shared genomic response to selection for monogamy across lineages (Young et al., 2019). These conserved molecular programs are thought to operate within the so-called ‘social decision-making network’ in vertebrates, and comprise interacting regulatory modules, including transcription factors and signaling pathways, that coordinate widespread transcriptional responses (Rittschof et al., 2014). These conserved gene regulatory/expression modules are often fine-tuned by species-specific regulatory changes which allow for both stable evolutionary shifts and the rapid behavioral plasticity required to navigate dynamic social environments (Alward et al., 2020; Renn et al., 2008).

In this study, we used cichlid fishes as a study system to investigate how the brain transcriptome is associated with both phylogenetic (i.e. inter-specific) and plastic (i.e. intra-specific) changes in social behavior. The adaptive radiation of cichlid fishes in Lake Tanganyika is the most diverse cichlid radiation and comprises approximately 240 species divided into 12 tribes that evolved in just about 10 million years (Ronco et al., 2021). It is characterized by extensive phenotypic diversity (Salzburger, 2018) including a range of social phenotypes that evolved repeatedly and independently (Sefc, 2011; Sorigue et al., 2026). In particular, the Lamprologini, the most species-rich tribe of the radiation, has attracted scientific attention for its diversity in social structures ranging from solitary individuals to complex hierarchical and cooperative societies (Lein & Jordan, 2021; Satoh et al., 2026). In the present study, we tested whether the evolution of sociality and social plasticity relies on the same molecular mechanisms. We investigated the molecular signatures underlying variation in sociality (i.e. solitary vs. social-living) at both the inter- and intra-specific levels across three closely related species, comprising two facultative social species (*Neolamprologus multifasciatus* and *Neolamprologus meeli*) and an obligately solitary species (*Lamprologus ornatipinnis*). To address these questions, we combined Differential Gene Expression (DGE) analyses with Gene Set Enrichment Analysis (GSEA) in two major brain regions that host most nodes of the social behavior network (Goodson, 2005; O’Connell & Hofmann, 2012): the telencephalon and the diencephalon. The neurogenomic signature of social plasticity was established by comparing the social and solitary behavioral phenotypes (aka ethotypes) within the facultative social species, whereas the neurogenomic signature for the evolution of social behavior (i.e. transition from solitary to social) was investigated by comparing the social ethotypes of the facultative social species with the obligatory solitary species. This framework allowed us to assess whether the neuromolecular mechanisms associated with the evolution of social living are shared across independent evolutionary transitions and whether these same mechanisms are also deployed flexibly within species to generate alternative social phenotypes.

## Materials and Methods

### Phenotypic characterization and ancestral state reconstruction of sociality across Lake Tanganyika cichlid species

Sociality was operationalized based on the observed social group size of each species in light of the Lake Tanganyika cichlid phylogeny (Ronco et al., 2021). We quantified social group size in an unsupervised manner using the raw observational data from Widmer et al. (2019a) collected as part of the study described in Widmer et al. (2019b).

In brief, in this study cameras were deployed underwater for a minimum of two hours, and images were captured during the second hour to avoid any disturbance created by the diver deploying the cameras. Still frames were extracted at 10-second intervals, from which observed species and numbers of individuals observed in each frame for each species was taken. For each deployment and species, we calculated the maximum number of individuals observed in a single frame (N_max_). Deployments with N_max_ = 0 were excluded from further analyses. We then computed the median N_max_ across all deployments for each species. Species with a median N_max_ of exactly 1 were classified as solitary, whereas species with a median N_max_ greater than 1 were classified as social-living (see Supplementary Table 1 for the median N_max_ obtained for each species). Information on the social phenotype (solitary vs. social-living) was available for 149 species.

Ancestral state reconstruction of sociality was computed by inferring the social trait (solitary vs. social-living) probabilities at all nodes of the Lake Tanganyika cichlid phylogeny (Ronco et al., 2021) for which individual counts were available. Maximum likelihood (ML) ancestral state reconstruction was conducted using the ace function implemented in the ape package (version 5.8; Paradis & Schliep, 2019) in R (version 4.5.2). Two models of discrete character evolution were compared: an equal-rates model (ER), which assumes a single transition rate between states, and an all-rates-different model (ARD), which allows different transition rates between solitary and social-living states. For each model, log-likelihood values were used to compute Akaike Information Criterion (AIC) scores. Model fit was assessed using ΔAIC and a likelihood ratio test (LRT) comparing the nested models. Neither ΔAIC (−0.29) nor the LRT (p = 0.13) supported a better fit of the more complex ARD model (Table 1). Therefore, the simpler ER model was retained for subsequent analyses. Marginal ancestral state probabilities at each internal node were then estimated under the ER model, providing the likelihood of each trait state at each node. To complement the ML approach, ancestral state reconstruction was also performed using a Bayesian framework implemented in MBASR (version v2023.10.27 ; Heritage, 2021). Analyses were conducted under a single-rate model of trait evolution for unordered discrete characters. Markov chain Monte Carlo (MCMC) sampling was run for 10,000 iterations to estimate posterior probabilities of trait states at each node. This approach integrates over parameter uncertainty to infer the posterior distribution of ancestral states. The output consisted of node-wise posterior probabilities for each trait state. The trait probabilities at all nodes are visualized in Supplementary Figure 1 (ML-based reconstruction) and Supplementary Figure 2 (Bayesian reconstruction), with details proved in Supplementary Table 2.

**Table 1.**
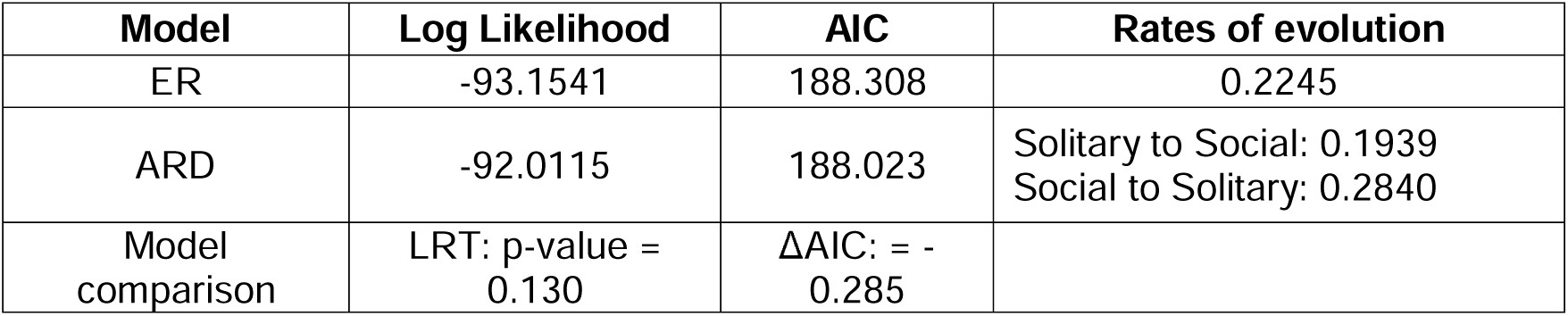
Model comparison for ancestral state reconstruction of group-living. Maximum likelihood models of discrete character evolution were compared using log-likelihood, Akaike Information Criterion (AIC), and likelihood ratio tests (LRT). The equal-rates model (ER) assumes a single transition rate between solitary and social-living states, whereas the all-rates-different model (ARD) allows distinct transition rates for each direction (solitary to social-living and social-living to solitary). ΔAIC was calculated as AIC_ARD_ – AIC_ER_. The likelihood ratio test compares the fit of the nested models. As neither ΔAIC nor the LRT (*p* = 0.130) supported a significantly better fit of the more complex ARD model, the ER model was retained for subsequent ancestral state reconstruction analyses.

### Selection of species

To investigate the evolution of sociality and its phenotypic plasticity, we selected three closely related species based on trait reconstruction, which allowed us to design experimental comparisons. Specifically, we included two facultative social species to investigate two independent instances of behavioral plasticity, and one obligately solitary species to contrast with each social species and thereby investigate two independent transitions toward sociality. To this end, we selected three species that have diverged in a very short period of time: *Neolamprologus multifasciatus*, *Neolamprologus meeli*, and *Lamprologus ornatipinnis*. Of the three, *N. multifasciatus* diverged shortly before *N. meeli* from a common ancestor, and *L. ornatipinnis* diverged from the common ancestor of all three species approximately 4 million years ago (Ronco et al., 2021). Under the single transition rate model, the trait reconstruction of the three species favored a solitary state, with 63.2% (ML) / 62.9% (Bayesian) probability at the root node and 64.4% (ML) / 64.0% (Bayesian) at the *N. meeli* - *L. ornatipinnis* divergence node (Figure 1). In this scenario, *N. multifasciatus* and *N. meeli* diverged from a most probable solitary ancestor, evolving social living independently. Additionally, the two social species (*N. multifasciatus* and *N. meeli*) are facultative social-living, with both solitary and social individuals occurring within populations. This intraspecific variation provides an opportunity to investigate behavioral plasticity and compare the mechanisms underlying social and solitary lifestyles both within and between species.

**Figure 1.**
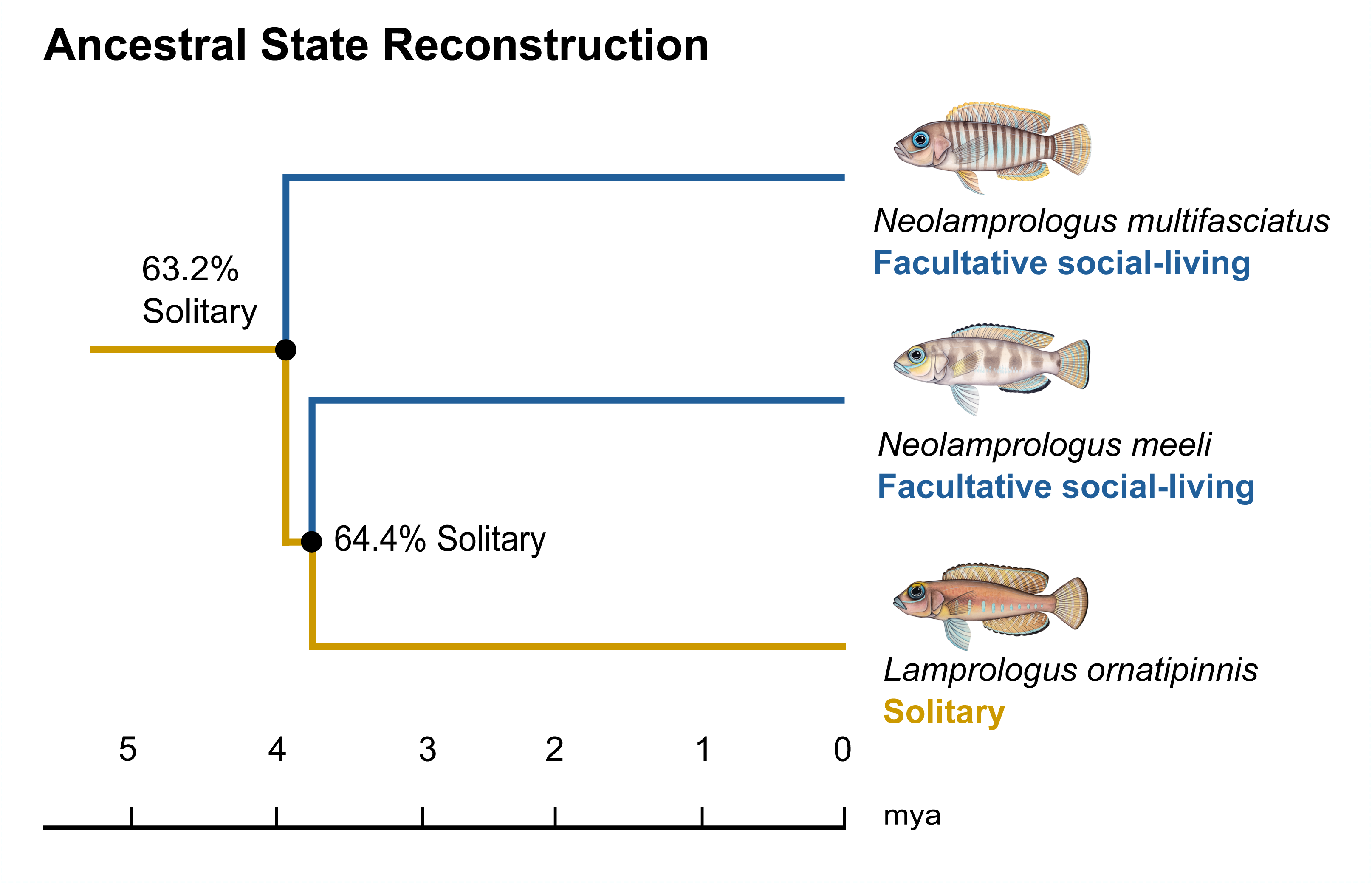
Ancestral state reconstruction and brain-region-specific gene expression differences in Lake Tanganyika cichlids. **(a)** Phylogenetic relationships among *Neolamprologus multifasciatus*, *Neolamprologus meeli*, and *Lamprologus ornatipinnis* with inferred ancestral states of social/solitary phenotypes. Colors indicate trait states (blue = facultative social-living, gold = solitary). Percentages at internal nodes correspond to the probability of a solitary ancestral state inferred under the equal-rates (ER) model using maximum likelihood. The time scale (in million years ago, mya) is shown along the x-axis and is taken from Ronco et al. (2021).

*N. multifasciatus* is an obligate shell-brooding cichlid endemic to Lake Tanganyika that lives on sandy bottoms where empty *Neothauma tanganyicense* shells serve as breeding chambers and refuges. *N. multifasciatus* typically forms social groups of four to five adults and multiple juveniles. Group members interact frequently, and intergroup interactions are common among close neighbors (Bose et al., 2022; Jordan et al., 2026). Occasionally, solitary, physically isolated individuals are observed with little to no social interaction. *Neolamprologus meeli*, another sandy-bottom shell-brooder, commonly lives in groups around shell nests. These groups are often embedded within a larger matrix of other *N. meeli* groups and different shell-brooding species like *N. brevis* and *N. temporalis* (Saeki et al., 2022; Sunobe & Munehara, 2003). Solitary *N. meeli* individuals can also be found in isolated patches of sand, where the presence of conspecifics and other fish is very limited. *Lamprologus ornatipinnis* males occupy territories alone and have minimal, if any, social interactions (Gordon & Bills, 1999).

### Collection of biological samples

Field work was carried out with approval from the Zambian Department of Fisheries, under study permits issued by the government of Zambia (No. SP260718/7-21 and SP297185/4-22) and in conjunction with memorandum of understanding MOU 101/14/11. The study species are listed as ‘least concern’ on the IUCN Red List of Threatened Species. All methods adhered to the ASAB/ABS Guidelines for the Use of Animals in Research.

Adult males of all the three study species were collected for biological sampling at Lake Tanganyika in October 2023 by scuba diving along the lake bottom near Chikonde island (8°42’48.6"S 31°07’26.6"E). For a detailed description of the sampling area see (Ma et al., 2025). Social-living individuals were selected from groups containing three or more fish. Solitary males were collected after ensuring that no other conspecific was occupying the same nest. A total of 30 individuals (six *L. ornatipinnis*, six social-living *N. multifasciatus*, six solitary *N.multifasciatus*, six social-living *N. meeli*, and six solitary *N.meeli*) were collected. Fish were captured and euthanized using an overdose of MS-222 (tricaine methanesulfonate, 1 g l^−1^), followed by rapid spinal cord transection. Brain dissections were performed within 10-15 minutes after capture, and whole-brain tissue was immediately stored in RNAlater (Sigma-Aldrich) and kept at 4°C until transportation to the laboratory. In the laboratory, samples were kept at -80°C. The telencephalon and diencephalon were dissected under a stereomicroscope (Zeiss; Stemi 2000) and transferred to 1 ml of QIAzol lysis reagent (RNeasy Lipid Tissue MiniKit, Qiagen). Four samples were excluded from further analyses due to poor tissue quality following RNAlater storage and subsequent dissection (two social-living *N. multifasciatus* individuals, one solitary *N. multifasciatus* individual, and one social-living *N. meeli* individual).

All methods are reported in accordance with ARRIVE guidelines (Percie Du Sert et al., 2020).

### RNA extraction and sequencing

Brain tissue was homogenized with a pestle in an Eppendorf tube on ice. RNA was then extracted using the RNeasy Lipid Tissue Mini Kit (Qiagen) following the manufacturer’s protocol (RNase-Free DNase set; Qiagen), with an additional DNAse cleaning step. RNA quality and concentration for each sample were determined using a NanoDrop 1000 spectrophotometer (Thermo Scientific). Samples were normalized to 200 ng/µl and processed following the MGIEasy Fast RNA Library Prep Set v1.0 protocol. The mRNAs were captured using the VAHTS mRNA Capture Beads v2.0 (Vazyme) and eluted in Low-TE buffer. Fragmentation and reverse transcription were performed, followed by single-strand synthesis and end-repair. PCR-mediated adaptor ligation was then carried out, followed by cleanup and PCR amplification to incorporate barcodes using the UDB PCR Primer Mix. The amplified products were purified, quantified using the Qubit dsDNA fluorescence assay (Thermo Fisher Scientific), and their size distributions (target length 200-400bp) was assessed using a Fragment Analyzer (Agilent Technologies). The purified PCR products were then circularized to generate DNA Nanoballs using the MGIEasy Dual Barcode Circularization kit. Library concentration and fragment-size distributions were assessed using Qubit quantification and a Fragment Analyzer (Agilent Technologies), respectively. Libraries were then sequenced on the MGI DNBSEQ-G400 using FCL SE 100, and the resulting sequence data were provided in FASTQ.

Raw sequencing reads were assessed for quality using FastQC (v0.12.1), and processed using Fastp (v0.23.4), which performs quality filtering by trimming bases with Phred scores below 15, removing reads with a high proportion of low-quality or ambiguous bases, and automatically detecting and trimming adapter sequences.

### Mapping and read quantification

FASTQ files were mapped to the Nile tilapia genome (*Oreochromis niloticus*; RefSeq accession GCF_001858045.2, female) using the STAR aligner (v2.7.11b; Dobin et al., 2013). The Nile tilapia was used as the reference genome due to its high-quality annotation and its phylogenetic equidistance to all cichlid species of the adaptive radiation of cichlid fishes in Lake Tanganyika (El Taher et al., 2020; Ronco et al., 2021). During mapping, up to 10 multimapping locations per read were allowed, and minimum match and score thresholds were set using *--outFilterMatchNminOverLread 0.4* and --*outFilterScoreMinOverLread 0.4*. Gene-level raw counts were obtained from the aligned BAM files using featureCounts (v2.0.2), assigning reads to genomic features based on the reference annotation.

### Outlier detection

We used variance-stabilized counts from the DESeq2 R package (v1.50.2; Love et al., 2014) to assess sample relationships and detect potential outliers. For each brain region, counts were first filtered to remove low-expression genes, following DESeq2’s recommendations. Principal component analysis (PCA) was then performed on the variance-stabilized transformed gene counts using the *prcomp()* function in R. Outlier detection was performed by calculating z-scores for the first four principal components within each group. Euclidean distances from the group centroid in standardized PC space were computed, and samples exceeding a distance threshold of 3 standard deviations were flagged as outliers. In doing so one sample of *L. ornatipinnis* was identified as outlier (3.7xSD) and excluded from subsequent analyses.

### Differential gene expression analysis

For intraspecific analysis, we compared the solitary and social-living individuals within each facultative social-living species (*N. multifasciatus* and *N.meeli*). Interspecific analyses compared the obligate solitary (*N. ornatipinnis*) with each ethotype of the facultative species – solitary and social-living. Therefore, we performed two intraspecific and four interspecific planned comparisons for each brain region. Pairwise comparisons were made using DESeq2 variance-stabilized counts with default parameters. Genes were considered differentially expressed with a fold-change of 1 between comparisons and an adjusted p-value of 0.05 after Benjamini–Hochberg correction for multiple testing.

### Functional enrichment analysis

Gene Set Enrichment Analysis (GSEA) was conducted for both Gene Ontology (GO) and Kyoto Encyclopedia of Genes and Genomes (KEGG) pathways using the *gseGO* and *gseKEGG* functions from the *clusterProfiler* package. Genes were ranked based on the ‘stat’ value derived from differential expression analysis in DESeq2, which reflects both effect size and variance, and the complete ranked gene list was used as input. GO enrichment analysis was performed for the Biological Process ontology using annotation data retrieved via *AnnotationHub* for *Oreochromis niloticus* (OrgDb: AH119811), and redundant GO terms were reduced using the *simplify* function to improve interpretability. Statistical significance was assessed using a Benjamini–Hochberg correction, and enriched gene sets were filtered using an adjusted p-value threshold of 0.05.

To identify biological processes associated with behavioral plasticity and shared between facultative species, GSEA results were further processed to detect common enriched GO terms and KEGG pathways across comparisons, namely *N. multifasciatus* (solitary vs social-living) and *N.meeli* (solitary vs social-living). To identify shared functional modules, semantic similarity analysis was performed using the *GOSemSim* package in R. GO terms from both comparisons were combined, and pairwise semantic similarity was computed using the Wang method based on the Biological Process ontology. The resulting similarity matrix was then used for hierarchical clustering (average linkage; distance = 1 − similarity), and clusters were defined by cutting the dendrogram at a fixed height (h = 0.75). GO terms were annotated by dataset of origin, and clusters containing terms from both datasets were retained as shared clusters. Shared KEGG pathways were identified by intersecting significant gene sets between the two species.

Similarly, we build a second similarity matrix with all categories enriched in the four interspecific comparisons, and hierarchical clustering and cutting were performed as described for the intraspecific scenario. The interspecific comparisons for each facultative species yielded enriched categories that were specific to: (i) the solitary phenotype, when the GO terms were only enriched in the *L. ornatipinnins* vs. the solitary individuals of the facultative species (ii) the social-living phenotype, when the GO terms were only enriched in the *L. ornatipinnins* vs. the social-living individuals of the facultative species; and (iii) both when the GO terms were enriched in the two comparisons. Because our primary focus was the evolution of social living, downstream analyses were restricted to clusters containing a majority (>70%) of terms specific to the social-living phenotype.

Clusters were named manually by assigning a category based on the biological functions represented by the GO terms within each cluster. The direction of the enrichment of each cluster (upregulated in solitary or in social-living phenotype) was considered upregulated when 70% of the categories had the same direction based on the Normalized Enrichment Score (NES). Otherwise, we considered “mixed” sign of enrichment.

## Results

### Phenotypic characterization of Lake Tanganyika cichlid species

The unsupervised quantification of social group size based on observational data (Widmer et al., 2019a) provided phenotypic information for 149 species. Of these, 89 species had a median maximum social group size (N_max_) value of 1 across all deployments and therefore we classified them as solitary, whereas 60 species had a median N_max_ > 1 and were classified as social-living (Figure 2). The dataset included observations of all three species of the study: *L. ornatipinnis* had a median N_max_ of 1, consistent with its solitary nature, while *N. multifasciatus* and *N. meeli* had a median N_max_ of 7 and 2, respectively (Figure 2; Supplementary Table 1).

**Figure 2.**
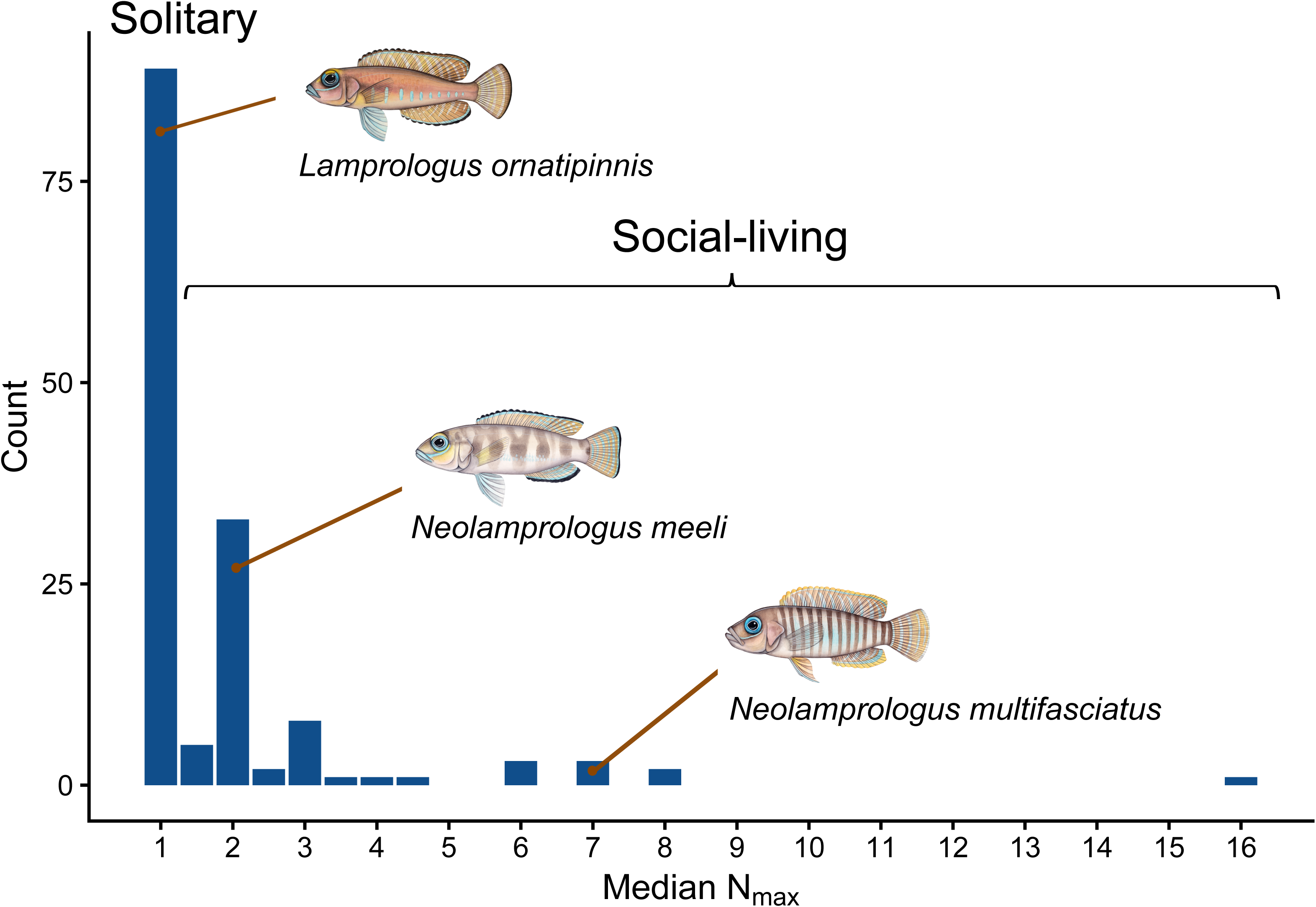
Distribution of the observed median maximum social group size across cichlid species. The histogram shows the distribution of median maximum group size (N_max_) across species for which information on social group size was available (Widmer et al., 2019a). The three focal species are indicated at their respective positions in the distribution. For the purposes of this study, species were classified as solitary when N_max_ = 1, and as social-living when N_max_ > 1.

### Sequencing and mapping performance

RNA-sequencing yielded a mean of ∼ 50 million reads per sample. On average, across samples, 89% of the reads were uniquely mapped to the reference genome and 3.2% mapped in multiple loci. The average read length across all samples was 94.85 base pairs per read. Per-sample mapping information is reported in Supplementary Table 3. The gene count matrix (Supplementary Data 1) comprised 42,646 annotated genomic features, including protein-coding genes as well as non-coding and other annotated *loci* (e.g., pseudogenes and regulatory elements). Of these, 35,218 had at least 1 count in at least one sample.

### Brain whole-transcriptome characterization

Principal component analysis (PCA) of brain gene expression profiles revealed that the primary axes of variation (PC1 and PC2) were largely driven by species identity in both brain regions (Figure 3). For the telencephalon transcriptome, *L. ornatipinnis* clustered closely with *N. meeli*, while *N. multifasciatus* was more separated, consistent with their phylogenetic relationships. In contrast, in the transcriptome of the diencephalon, individuals clustered by species, with both facultative species ( *N. meeli*, *N. multifasciatus*) and *L. ornatipinnis* forming well-separated groups. Across both brain regions, phenotypic differences (social-living vs solitary) contributed comparatively little to the overall variance captured by the first two principal components, as evidenced by the substantial overlap between phenotypes within species (Figure 3) when considering the whole transcriptome.

**Figure 3.**
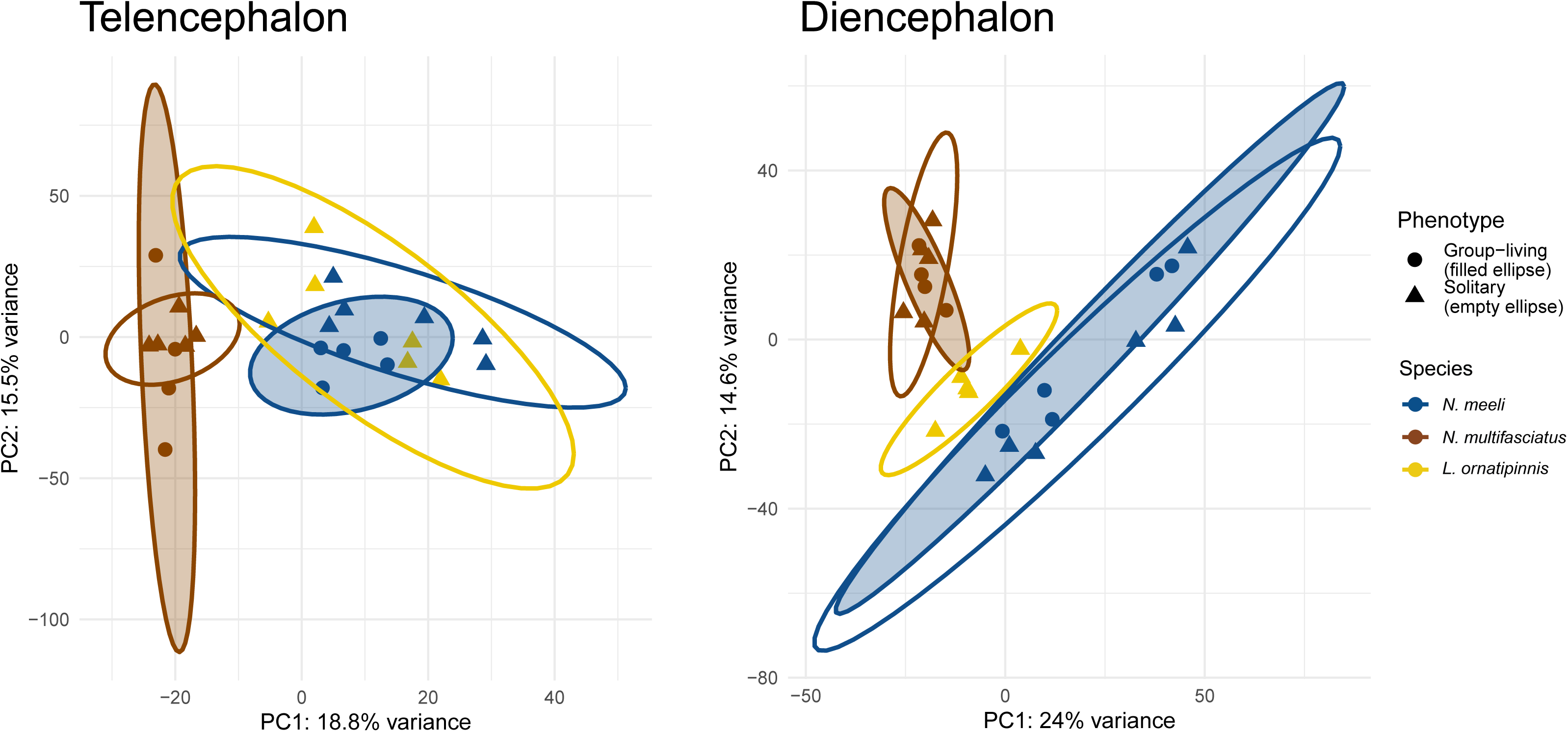
Principal component analysis of gene expression across brain regions, species, and phenotypes. Principal component analysis (PCA) of transcriptomic profiles in the diencephalon (left) and telencephalon (right). Each point represents an individual sample, colored by species (*N. meeli*, *N. multifasciatus*, and *L. ornatipinnis*) and shaped by phenotype (circles: group-living; triangles: solitary). The first two principal components (PC1 and PC2) are shown, with the percentage of variance explained indicated on each axis. Ellipses represent the 95% confidence intervals for each species group; filled ellipses correspond to social-living individuals, and open ellipses correspond to solitary individuals. Separation along principal components reflects mainly variation associated with species.

### Differential Gene Expression and Functional Enrichment in relation to intra-specific variation in sociality

Using thresholds of log fold change of 1 and adjusted p-value (FDR) ≤ 0.05, intraspecific differential gene expression analyses between the social-living and solitary individuals revealed 8 differentially expressed genes (DEG) in *N. multifasciatus* and 20 DEG in *N. meeli* in the telencephalon, and 9 DEGs in *N. multifasciatus* and 64 in *N. meeli* in the diencephalon (Figure 4A; Supplementary Tables 4 and 5). No DEGs were shared between species within the same brain region. In contrast, several genes were consistently differentially expressed across brain regions within species (see Supplementary Data 2 for the complete list of genes after differential gene expression analyses of the telencephalon, and Supplementary Data 3 for the complete list of genes after differential gene expression analyses of the diencephalon).

**Figure 4.**
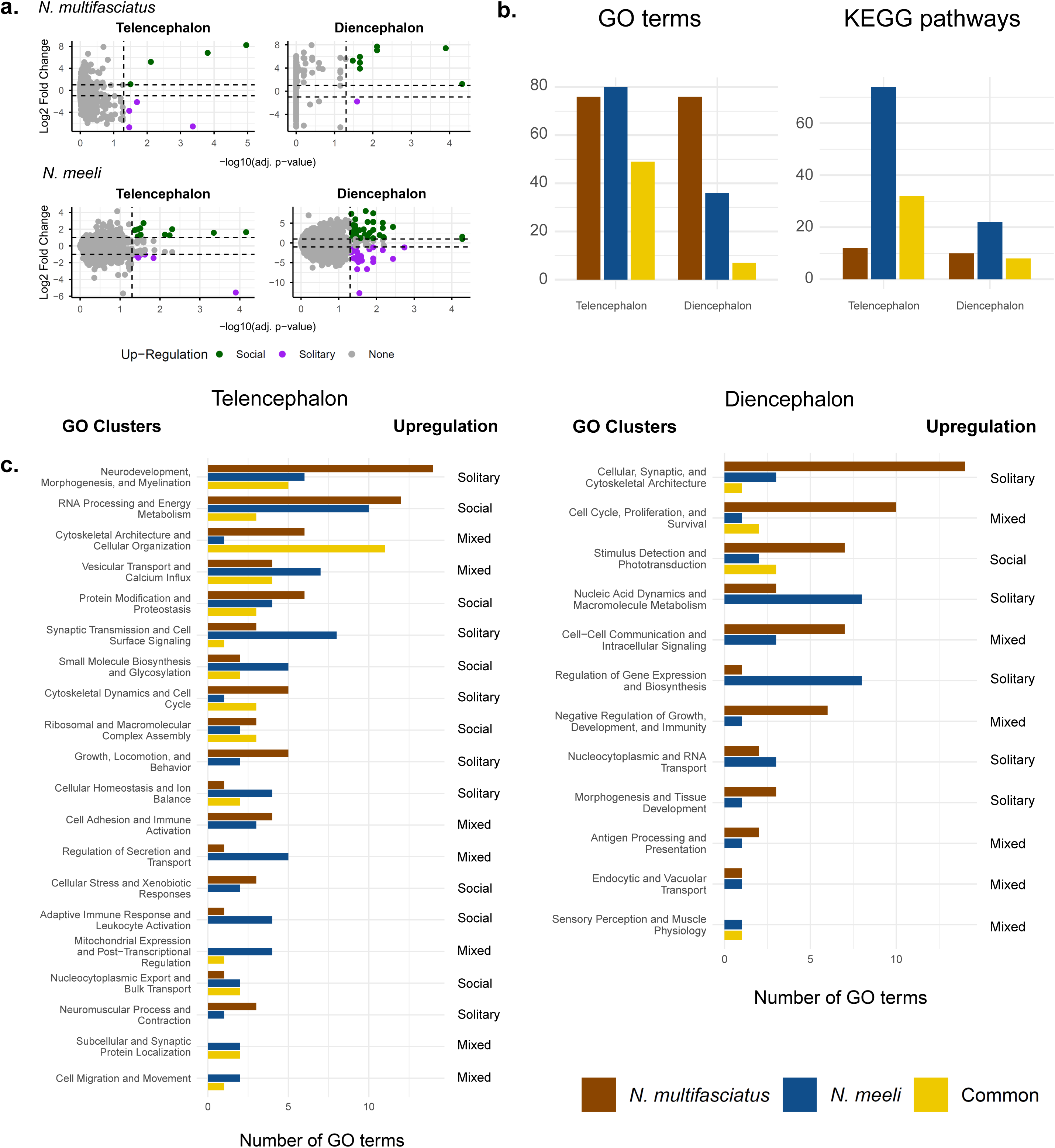
Intraspecies differential gene expression and functional enrichment of differentially expressed genes associated with social phenotypes across brain regions. (a) Intraspecific differential gene expression between social-living and solitary individuals within *N. multifasciatus* (top) and *N. meeli* (bottom) in the telencephalon and diencephalon. Each point represents a gene. The x-axis shows statistical significance (−log10 adjusted *p*-value), and the y-axis shows expression difference (log2 fold change). Green points indicate genes upregulated in social individuals, purple points indicate genes upregulated in solitary individuals, and gray points indicate non-significant genes. Dashed vertical and horizontal lines denote significance (p-value = 0.05) and fold-change (logFC = 1) thresholds, respectively. Sample sizes for each group are as follows: telencephalon: *N. multifasciatus* solitary (N = 5), *N. multifasciatus* social-living (N = 4), *N. meeli* solitary (N = 6), *N. meeli* social-living (N = 5); diencephalon: *N. multifasciatus* solitary (N = 5), *N. multifasciatus* social-living (N = 4), *N. meeli* solitary (N = 6), *N. meeli* social-living (N = 5). **(b)** Summary of enriched functional categories identified among differentially expressed genes in the telencephalon and diencephalon of *N. multifasciatus* and *N. meeli*. Bar plots show the number of significantly enriched GO terms and KEGG pathways detected in each species, as well as those shared between species (“Common”). **(c)** Distribution of enriched GO-term clusters in the diencephalon (left) and telencephalon (right). Bars indicate the number of enriched GO terms assigned to each functional cluster for *N. multifasciatus* (brown), *N. meeli* (blue), or shared between species (gold). Clusters are ordered by number of enriched GO terms. The column “Upregulation” indicates whether genes contributing to enrichment were predominantly upregulated in solitary individuals, social-living individuals, or showed a mixed pattern of regulation. Functional cluster names summarize broad biological processes represented by the enriched GO terms. The x-axis represents the number of GO terms assigned to each cluster. *Abbreviations:* GO, Gene Ontology; KEGG, Kyoto Encyclopedia of Genes and Genomes.

Comparative Gene Set Enrichment Analysis revealed distinct functional landscapes for the telencephalon and diencephalon.

#### Telencephalon

In the telencephalon, 125 GO terms were enriched between the two social phenotypes in *N. multifasciatus* and 129 in *N. meeli*, of which 49 were shared between species (Figure 4B). Clustering analysis identified 32 functional clusters, 20 of which contained GO terms from both species, while 15 included at least one GO term shared between species (Supplementary Table 6). The 20 shared clusters encompassed 91.4% (192/210) of all enriched GO terms (Supplementary Figure 3; Supplementary Data 4), indicating substantial convergence in the biological processes associated with social phenotype across both facultative species. Shared GO terms and functional clusters were enriched in the same direction across both species, matching their respective social phenotypes (social-living or solitary), demonstrating a conserved functional response to the social environment.

The telencephalon of social-living individuals in both species exhibited a robust, conserved upregulation of translational and energetic machinery. Within the clusters *Protein Modification and Proteostasis*, and *Ribosomal and Macromolecular Complex Assembly*, the GO terms *translation* (p-value < 10e-7), *protein maturation* (p-value < 10e-6), and *ribosome biogenesis* (p-value < 10e-7) were significantly enriched in both species, together with KEGG *Ribosome* enrichment (p-value < 10e-4). Similarly, the large cluster *RNA Processing and Energy Metabolism* was consistently upregulated in social-living individuals and included several catabolic pathways associated with lipid degradation and the generation of precursor metabolites and energy (Figure 4C). These patterns were supported by enrichment of KEGG pathways including *Oxidative phosphorylation* (*N. multifasciatus* p-value = 2.5e-9; *N. meeli* p-value = 2.1e-3), *Fatty acid degradation* (p-value < 5e-2), and *Valine, leucine and isoleucine degradation* (p-value < 10e-5) (Supplementary Table 6).

In contrast, solitary individuals of both species show upregulated neurodevelopmental and structural functions in the telencephalon. The large cluster *Neurodevelopment, Morphogenesis, and Myelination* was consistently upregulated and included GO terms related to tissue development and cell differentiation, such as *plasma membrane bounded cell projection morphogenesis* (p-value < 10e-4) and *regulation of developmental process* (p-value < 5e-2). Additional species-specific upregulated GO terms in solitary individuals included *neuron differentiation* (*N. multifasciatus* p-value = 1.5e-8), *axon guidance* (*N. multifasciatus* p-value = 3.3e-8), *presynapse assembly* (*N. meeli* p-value = 4.6e-3), and *regulation of neurogenesis* (*N. meeli* p-value = 1.8e-2). The cluster *Cytoskeletal Dynamics and Cell Cycle* further supported patterns of neuronal differentiation and tissue remodeling, whereas the cluster *Synaptic Transmission and Cell Surface Signaling* highlighted processes associated with neural development and synaptic plasticity (Figure 4C). Consistent with these findings, several KEGG pathways associated with structural remodeling and neuronal activity were enriched in solitary individuals, including *Extracellular Matrix-receptor interaction* (p-value *<* 0.05), *Focal adhesion* (p-value < 10e-3), and *Calcium signaling pathway* (p-value < 10e-2) (Supplementary Table 6).

Two clusters contained GO terms upregulated in both social phenotypes (solitary and social-living). In the *Vesicular Transport and Calcium Influx* cluster, social-living individuals upregulated Golgi and hormone transport, while solitary individuals upregulated calcium transport and synaptic vesicle recycling. In the cluster *Cytoskeletal Architecture and Cellular Organization*, social-living individuals upregulated *vesicle trafficking*, *mitochondrial function*, and *lysosome biogenesis*. Conversely, solitary individuals upregulated *cytoskeletal, cell junction, and extracellular matrix organization*, reinforcing patterns of structural remodeling (Supplementary Table 6).

#### Diencephalon

In the diencephalon, 83 GO terms were enriched in *N. multifasciatus* and 41 in *N. meeli* when comparing solitary vs. social-living individuals. Of these 7 were shared between species (Figure 4B). Clustering analysis identified 20 functional clusters, 12 of which contained GO terms from both species, while 4 included at least one GO term shared between species (Figure 4C; Supplementary Table 6). The 20 shared clusters encompassed 82.1% (96/117) of all enriched GO terms (Supplementary Figure 3; Supplementary Data 4). Unlike the telencephalon, overlap in enriched GO terms between species was limited. Moreover, enrichment direction was frequently inconsistent between species, even within the same functional clusters. Similarly, six of the eight shared KEGG pathways exhibited opposite enrichment directions, indicating upregulation in different social phenotypes across species (Supplementary Table 6).

Among the clusters containing shared GO terms, *Stimulus Detection and Phototransduction* was mostly upregulated in social living individuals. However, two shared categories of the cluster had opposite signs between the species: *phototransduction* and *detection of light stimulus* were significantly enriched in social-living *N. multifasciatus* (NES ≥ 2.28, p-value < 10e4) but in solitary *N. meeli* (NES ≤ -1.88, p-value < 0.05). The same pattern was observed for *visual perception* in another cluster: *N. multifasciatus* NES = 2.6 (p-value = 1.2e-8) and *N. meeli* NES = -2.24 (p-value = 1.2e-4). Consistent with these findings, the KEGG pathway *phototransduction* was strongly enriched in both species (p-value = 1.78e-8) but with opposite enrichment directions (*N. multifasciatus*: 2.98, *N. meeli*: -2.7). Despite these differences, conserved patterns remained in upregulation of cell cycle in solitary individuals with shared categories as *cell cycle process* (p-value < 0.05) and *chromosome segregation* (p-value < 0.05), and supported by conserved upregulation of KEGG *cell cycle* (p-value < 0.05) and *ATP-dependent chromatin remodeling* (p-value < 0.05).

The complete list of enriched GO and KEGG terms in the intraspecies analysis can be consulted in Supplementary Data 5.

### Differential Gene Expression and Functional Enrichment in relation to inter-specific variation in sociality

Interspecific differential expression analyses were conducted between the obligately solitary species *L. ornatipinnis* and the two groups – solitary and social-living – of the facultative social species: The rational for these comparisons is that, whereas differences between solitary individuals of the different species should reflect phylogenetic divergence, differences between the mandatory solitary *L. ornatipinnis* and the social-living ethotypes of the facultative social-living species should be associated with the evolution of social-living. In the telencephalon, 1,293 and 1,353 DEG were identified relative to solitary and social-living *N. multifasciatus*, and 929 and 1,137 DEG relative to solitary and social-living *N. meeli*, respectively. In the diencephalon, the corresponding comparisons yielded 898 and 719 DEG in *N. multifasciatus* solitary and social-living respectively, and 1,063 and 955 DEG in *N. meeli* (Figure 5A).

**Figure 5.**
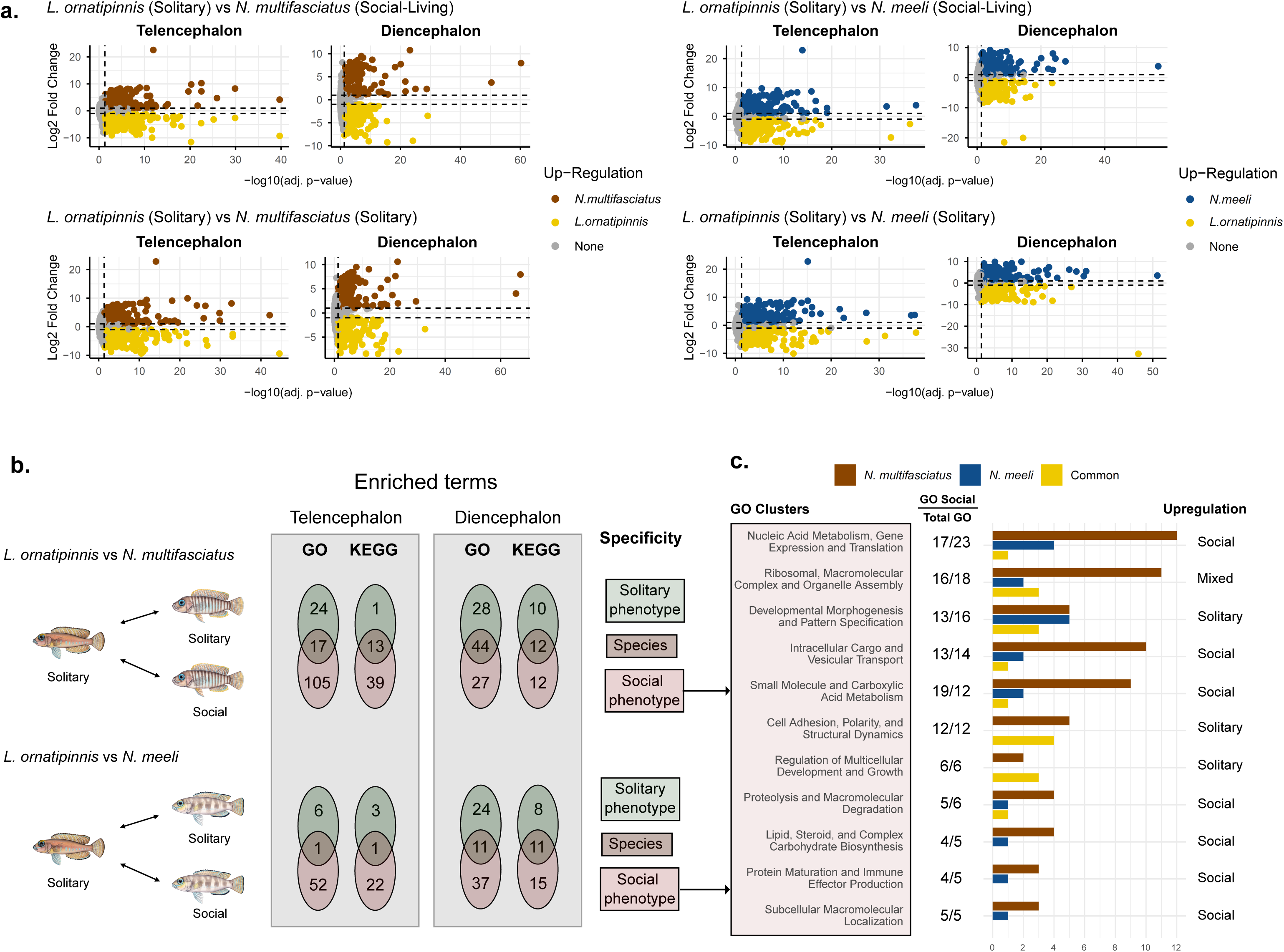
Interspecies differential gene expression and functional enrichment of differentially expressed genes associated with social phenotypes across brain regions. (a) Interspecific differential gene expression comparisons between the solitary species *L. ornatipinnis* and the facultatively social species *N. multifasciatus* (left) and *N. meeli* (right). Comparisons are presented using either social-living or solitary individuals from the facultatively social species. Colored points represent significantly differentially expressed genes, with colors indicating the species in which expression is higher; gray points indicate non-significant genes. Dashed vertical and horizontal lines denote significance (p-value = 0.05) and fold-change (logFC = 1) thresholds, respectively.. Sample sizes for each group are as follows: telencephalon: *N. multifasciatus* solitary (N = 5), *N. multifasciatus* social-living (N = 4), *N. meeli* solitary (N = 6), *N. meeli* social-living (N = 5), *L. ornatipinnis* solitary (N = 6); diencephalon: *N. multifasciatus* solitary (N = 5), *N. multifasciatus* social-living (N = 4), *N. meeli* solitary (N = 6), *N. meeli* social-living (N = 5), *L. ornatipinnis* solitary (N = 5).**(b)** Summary of GO and KEGG pathway enrichment identified from interspecies differential gene expression analyses comparing the solitary species *L. ornatipinnis* with *N. multifasciatus* and *N. meeli*. Comparisons were performed between *L. ornatipinnis* and both the solitary and social phenotypes of each facultatively social species. Venn diagrams show the number of enriched GO terms and KEGG pathways that were specific to the solitary phenotype, specific to the social phenotype, or shared between phenotypes in the telencephalon and diencephalon. **(c)** Functional clustering of GO terms uniquely enriched in the social phenotype for interspecies comparisons in the telencephalon. The left panel lists the major GO-term clusters, while the adjacent column indicates the number of GO terms assigned to each cluster relative to the total number of GO terms within that cluster (GO Social/Total GO). Bar plots show the contribution of *N. multifasciatus* (brown), *N. meeli* (blue), and GO terms shared between species (yellow) within each functional cluster. Clusters are ordered by number of enriched GO terms. The "Upregulation" column indicates whether enriched genes within each cluster were predominantly upregulated in social individuals, solitary individuals, or showed a mixed pattern of regulation. *Abbreviations:* GO, Gene Ontology; KEGG, Kyoto Encyclopedia of Genes and Genomes.

#### Telencephalon

Gene Set Enrichment Analysis in the telencephalon revealed a very strong phenotypic signal in both species by comparing the social ethotype of each species vs. the obligate solitary *L. ornatipinnis* While 41 GO terms were enriched between *L. ornatipinnis* and the solitary ethotype in *N. multifasciatus*, 122 showed significant enrichment between *L. ornatipinnis* and the social *N. multifasciatus*. Of these two sets, 17 GO terms were in common, reflecting phylogenetic rather than phenotype signal. Similarly, 7 GO terms were enriched between *L. ornatipinnis* and solitary *N. meeli*, but 53 were enriched when compared to the social phenotype, and 1 was in common. Similarly, KEGG pathway analysis also revealed a very strong effect of the phenotype. Combining both species’ comparisons, 61 terms were enriched in social-living, while 4 were enriched in solitary and 14 in both (; Supplementary Tables 7 and 8). This analysis identified GO and KEGG terms reflecting functional specificity to a single phenotype, as well as species-shared terms enriched in both phenotypes (Figure 5B). To further analyze the evolution of social living, we restricted our downstream analyses to the terms enriched exclusively in the social-living phenotype, as these reflect the evolution of social-living from the solitary ancestor.

Gene Set Enrichment Analysis in the telencephalon revealed a coordinated upregulation of core cellular processes in social-living individuals (Figure 5C). The large cluster *Nucleic Acid Metabolism, Gene Expression, and Translation* was highly upregulated, characterized by elevated DNA replication and repair pathways, *translation* (p-value < 10e-2), and extensive protein synthesis components (Supplementary Table 7). The cluster *Intracellular Cargo and Vesicular Transport* was also upregulated, showing enhanced nuclear-cytoplasmic export of mRNA and ribosomal subunits, *proton transmembrane transport* (p-value < 0.05), *Golgi vesicle transport* (*N. multifasciatus* p-value = 4.7e-4), and targeted protein localization to the mitochondria. Finally, the cluster *Small Molecule and Carboxylic Acid Metabolism* was significantly enriched in social-living individuals, alongside *2-Oxocarboxylic acid metabolism* KEGG pathway enrichment in solitary individuals of both species (p-value < 10e-2; Supplementary Table 8).

In contrast, solitary individuals showed enrichment of pathways associated with morphological specialization and neural wiring (Figure 5C). The first major functional category, *Developmental Morphogenesis and Pattern Specification*, was heavily enriched, characterized by pathways governing anatomical structure formation, tissue morphogenesis, and circulatory system development (Supplementary Table 7). Within this category we found representative terms such as *cell morphogenesis* (p-value < 0.05), *generation of neurons* (*N. meeli* p-value = 0.01), and *axon guidance* (*N. multifasciatus* p-value = 7.1e-4). This was tightly coupled with a second major cluster, *Cell Adhesion, Polarity, and Structural Dynamics*, marked by an upregulation of terms such as *cell-cell adhesion* (p-value < 10e-3), *establishment or maintenance of cell polarity* (p-value < 0.05), and *protein folding* (p-value < 10e-4). In this line, KEGG pathway *Extracellular Matrix-receptor interaction* (p-value < 10e-3) was upregulated in solitary individuals in both species (Supplementary Table 8).

Finally, the cluster *Ribosomal, Macromolecular Complex and Organelle Assembly* had terms upregulated in both phenotypes (Figure 5C). These included highly enriched mitochondria and ribosome assembly-related categories upregulated in social-living individuals, and terms associated with tissue development and cellular organization upregulated in solitary individuals (Supplementary Table 7).

#### Diencephalon

In contrast to the telencephalon, Gene Set Enrichment Analysis (GSEA) across species and social phenotypes of the diencephalon did not reveal a clear phenotypic signal. Between *L. ornatipinnis* and the solitary phenotype of *N. multifasciatus*, 28 GO terms were significantly enriched, compared with 27 GO terms between *L. ornatipinnis* and the social phenotype. 44 enriched GO terms were shared between the two comparisons. Similarly, 24 GO terms were enriched between *L. ornatipinnis* and the solitary phenotype of *N. meeli*, whereas 37 were enriched in the comparison with the social phenotype, and 11 GO terms were shared between both contrasts. KEGG pathway analysis likewise did not show a strong phenotypic signal: across both species, 18 enriched GO terms were unique to the solitary phenotype, 27 were unique to the social-living phenotype, and 23 were shared between both phenotypes. (Figure 5C; Supplementary Table 7).

Notably, no social-living-specific GO term was found in common between the two comparisons (*L. ornatipinnis* vs. social *N. multifasciatus* and *L. ornatipinnis* vs social *N. meeli*). Furthermore, the criteria used to assign clusters specific to social-living phenotype did not provide any social-living-specific cluster, as no cluster had more than 70% of terms only enriched in the social-living comparison (Supplementary Table 7). KEGG enrichment specific to the social living phenotype only showed one enriched pathway in common between both species: *Extracellular matrix-receptor interaction*, upregulated in solitary individuals (*N. multifasciatus* p-value = 1.7e-4, *N. meeli* p-value = 2.1e-2; Supplementary Table 8).

The complete list of enriched GO and KEGG terms in the intraspecies analysis can be consulted in Supplementary Data 5.

### Common brain transcriptome signatures of behavior plasticity and evolution

To identify shared brain gene expression signatures driving behavioral plasticity and the evolution of sociality, we crossed the enriched functional terms from our intra- and interspecific analyses. We focused on terms commonly enriched during phenotypic transitions within both facultative species, as well as those shared between the social ethotypes of both facultative species and the obligate solitary lineage (social-living-specific terms).

In the telencephalon, this approach revealed a robust upregulation in social-living phenotypes (positive NES) of core metabolic and protein-processing pathways vital for energetic homeostasis, including translation, protein folding, and mitochondrial respiratory chain complex assembly (Table 2). Conversely, processes governing structural dynamics and tissue remodeling were upregulated in social-living phenotypes (negative NES). Complementing the GO analysis, KEGG pathway enrichment further highlighted a shared metabolic and structural re-regulation spanning both behavioral plasticity and evolution. Social-living phenotypes exhibited a highly significant upregulation (positive NES) of pathways involved in protein turnover, degradation, and cellular energetics, while structural pathways linked to cellular adhesion and tissue scaffolding were consistently upregulated in solitary individuals (negative NES) across both intra- and interspecific scales (Table 2). No common GO nor KEGG term was found in common between all four comparisons in the diencephalon.

**Table 2.** Shared functional genomic signatures of behavioral plasticity and evolution. GO and KEGG terms significantly enriched across both intraspecies comparisons (behavioral plasticity) and interspecies comparisons (behavioral evolution) in *Neolamprologus multifasciatus* and *Neolamprologus meeli*. Behavioral plasticity captures differential gene expression between distinct ethotypes within a single species, while behavioral evolution captures differences between social-living and the obligate solitary *L. ornatipinnis*. NES: Normalized Enrichment Score; positive values indicate upregulation in social-living phenotypes, while negative values indicate downregulation in social-living phenotypes relative to solitary counterparts. Statistical significance is reported via p-values, with all included terms meeting the threshold for significant enrichment across both axes of comparison (p-value < 0.05). Cells highlighted in gray represent terms upregulated in solitary individuals (negative NES) while white cells are terms upregulated in social-living individuals (positive NES).

|  |  | **Behavior plasticity** |  | **Behavior evolution** |  |
|  |  |  |  |  |  |
|---|---|---|---|---|---|
| **GO term** | **Species** | **NES** | **p-value** | **NES** | **p-value** |
| Translation | *N. multifasciatus* | 2.69 | 1.47E-08 | 2.13 | 1.38E-08 |
|  | *N. meeli* | 2.05 | 2.05E-08 | 1.53 | 4.75E-03 |
| Protein folding | *N. multifasciatus* | 2.3 | 7.37E-08 | 2.11 | 3.95E-06 |
|  | *N. meeli* | 2.45 | 2.05E-08 | 2.09 | 3.35E-05 |
| Mitochondrial respiratory chain complex assembly | *N. multifasciatus* | 2.38 | 1.84E-06 | 2.19 | 1.43E-04 |
|  | *N. meeli* | 1.83 | 6.10E-03 | 2.02 | 2.48E-03 |
| Modification-dependent macromolecule catabolic process | *N. multifasciatus* | 1.56 | 2.00E-03 | 1.55 | 3.25E-03 |
|  | *N. meeli* | 1.46 | 1.21E-02 | 1.38 | 4.63E-02 |
| Mitochondrion organization | *N. multifasciatus* | 2.28 | 1.51E-08 | 2.35 | 1.38E-08 |
|  | *N. meeli* | 1.84 | 4.70E-04 | 1.95 | 2.96E-04 |
| Cell morphogenesis | *N. multifasciatus* | -1.83 | 1.08E-05 | -1.47 | 1.17E-02 |
|  | *N. meeli* | -1.9 | 5.34E-07 | -1.44 | 4.63E-02 |
| Plasma membrane bounded cell projection morphogenesis | *N. multifasciatus* | -1.91 | 2.99E-06 | -1.51 | 1.95E-02 |
|  | *N. meeli* | -1.84 | 1.40E-05 | -1.49 | 2.81E-02 |
| Regulation of developmental process | *N. multifasciatus* | -1.42 | 1.36E-02 | -1.52 | 8.65E-03 |
|  | *N. meeli* | -1.68 | 1.92E-04 | -1.66 | 1.26E-03 |
| Cell junction organization | *N. multifasciatus* | -1.74 | 3.85E-04 | -1.49 | 2.01E-02 |
|  | *N. meeli* | -1.89 | 7.69E-06 | -1.64 | 4.93E-03 |
| Establishment or maintenance of epithelial cell apical/basal polarity | *N. multifasciatus* | -1.76 | 3.40E-02 | -1.94 | 1.10E-02 |
|  | *N. meeli* | -1.85 | 1.83E-02 | -1.76 | 3.92E-02 |
| Regulation of cell differentiation | *N. multifasciatus* | -1.47 | 4.03E-02 | -1.67 | 3.25E-03 |
|  | *N. meeli* | -1.59 | 5.67E-03 | -1.68 | 4.10E-03 |
| **KEGG term** | **Species** | **NES** | **p-value** | **NES** | **p-value** |
| Proteasome | *N. multifasciatus* | 2.13 | 2.38E-04 | 2.45 | 4.93E-07 |
|  | *N. meeli* | 2.49 | 2.82E-06 | 1.93 | 2.93E-03 |
| Protein processing in endoplasmic reticulum | *N. multifasciatus* | 1.50 | 9.65E-03 | 1.69 | 6.26E-04 |
|  | *N. meeli* | 1.77 | 1.88E-04 | 1.91 | 3.71E-05 |
| Citrate cycle (TCA cycle) | *N. multifasciatus* | 1.90 | 7.91E-03 | 2.37 | 2.77E-05 |
|  | *N. meeli* | 1.90 | 3.41E-03 | 1.92 | 1.17E-02 |
| Mitophagy - animal | *N. multifasciatus* | 1.62 | 6.59E-03 | 1.56 | 7.02E-03 |
|  | *N. meeli* | 1.54 | 9.19E-03 | 1.63 | 7.13E-03 |
| Biosynthesis of cofactors | *N. multifasciatus* | 1.62 | 4.60E-03 | 1.41 | 3.98E-02 |
|  | *N. meeli* | 1.42 | 3.07E-02 | 1.48 | 1.52E-02 |
| Extracellular matrix-receptor interaction | *N. multifasciatus* | -2.64 | 2.54E-09 | -1.98 | 1.37E-04 |
|  | *N. meeli* | -1.43 | 3.63E-02 | -1.82 | 1.88E-03 |
| Cytoskeleton in muscle cells | *N. multifasciatus* | -2.60 | 2.54E-09 | -1.89 | 3.39E-06 |
|  | *N. meeli* | -1.65 | 5.65E-04 | -1.47 | 1.18E-02 |
| Cadherin signaling | *N. multifasciatus* | -1.44 | 3.82E-03 | -1.46 | 4.83E-03 |
|  | *N. meeli* | -1.56 | 5.65E-04 | -1.32 | 2.62E-02 |

## Discussion

Independent transitions to social living offer a natural experiment to examine whether or not evolution repeatedly recruits the same molecular machinery to build similar phenotypes. Here, we examined two independent transitions from solitary to social living within the lamprologine cichlids of Lake Tanganyika and asked whether or not the same neurogenomic programs are associated with both the evolution of sociality and the expression of alternative social phenotypes within species. Our results reveal a striking convergence in gene expression signatures in the telencephalon, where both evolutionary and plastic transitions were characterized by highly similar functional transcriptional signatures despite limited overlap at the level of individual differentially expressed genes. In contrast, the diencephalon exhibited comparatively weak and inconsistent transcriptional responses. Together, these findings suggest that repeated evolution of social living in cichlids involves the recurrent recruitment of conserved transcriptional programs in the telencephalon and provide support for the existence of a conserved behavioral genetic toolkit underlying social evolution and social plasticity.

One of the clearest patterns emerging from our analyses is the strong regional specificity of the neurogenomic response to social-living. Across both intra- and interspecific comparisons, the telencephalon consistently exhibited greater functional convergence than the diencephalon in response to social-living. Shared functional clusters encompassed most enriched GO terms in both facultative social species, enrichment directions were remarkably consistent, and interspecific comparisons yielded substantially more phenotype-specific than species-specific functional signatures in gene expression. In contrast, the diencephalon showed limited overlap of functional categories between species, frequent reversals in enrichment direction, and no functional clusters uniquely associated with social living. These findings are consistent with the established role of the telencephalon as a major integrative center for social cognition, learning, memory, and behavioral flexibility within the Social Decision-Making Network in teleost fish (Li et al., 2025; O’Connell & Hofmann, 2012; Teles et al., 2016). Rather than suggesting that the diencephalon is less involved in social behavior, our results indicate that conserved transcriptomic signatures distinguishing social phenotypes are concentrated in higher-order integrative regions, whereas hypothalamic and preoptic circuits may rely on more species-specific regulatory mechanisms or finer spatial patterns that are difficult to detect using bulk tissue transcriptomics.

Despite identifying relatively few differentially expressed genes within species, GSEA uncovered extensive functional reorganization associated with behavioral plasticity. When comparing solitary and social-living individuals within each facultative social-living species, only eight and twenty genes were differentially expressed in the telencephalon of *N. multifasciatus* and *N. meeli*, respectively, yet more than one hundred GO terms were significantly enriched in each species, with nearly half shared between them. This apparent discrepancy suggests a highly polygenic nature of social behavior. Rather than depending on a small number of genes with large transcriptional changes, alternative social phenotypes appear to arise through subtle but coordinated shifts across numerous genes belonging to common biological pathways. Similar patterns of regulation have been reported in other social systems and are consistent with theoretical expectations that complex behavioral traits emerge from changes in gene regulatory networks rather than from discrete "social genes" (Cole, 2014; Kapheim et al., 2015; Renn et al., 2016).Remarkably, highly overlapping functional programs characterized both behavioral plasticity and evolutionary divergence in the telencephalon. Across both analytical scales, social-living individuals consistently upregulated pathways associated with translation, protein folding, mitochondrial function, and protein catabolism. For example, *translation* and *mitochondrial respiratory chain complex assembly* were positively enriched at both scales in both species, and this metabolic signature extended to KEGG pathways, with *Proteasome* and *Citrate cycle (TCA cycle)* consistently upregulated in social-living individuals. Conversely, solitary individuals across species exhibited enrichment of pathways involved in cell morphogenesis, developmental regulation, cell junction organization, and cytoskeletal/adhesion dynamics – including *cell morphogenesis* and *Regulation of cell differentiation* – mirrored at the pathway level by consistent upregulation in solitary of *Extracellular matrix-receptor interaction* and *Cadherin signaling* across both species and analytical scales. The recurrence of these opposing transcriptional programs across independent evolutionary transitions and behavioral plasticity states suggest that the same ancestral molecular modules have been repeatedly recruited to generate social phenotypes.

Taken together, our results point to a conserved transcriptional trade-off associated with social behavior. Social living was consistently associated with transcriptional programs likely required to sustain the high metabolic demands imposed by continuous social interactions and behavioral flexibility. Processing social information requires the integration of multiple sensory inputs, the evaluation of social relationships, and the rapid adjustment of behavioral responses, all of which are expected to increase neuronal activity and biosynthetic demand. The high energetic investment observed in the telencephalon of social ethotypes supports the ’expensive brain’ framework of social energetics, where processing, predicting, and responding to complex conspecific cues shifts the baseline neural metabolic budget toward high-throughput synaptic activity (Coto & Traniello, 2022). Because maintaining group cohesion forces individuals to dynamically align their behavior with conspecifics, sociality is inherently linked to underlying physiological and metabolic constraints (Seebacher & Krause, 2017). In contrast, solitary individuals consistently exhibited in their gene expression profiles enrichment of pathways classically associated with neurodevelopmental processes, suggesting greater investment in maintaining or reorganizing neural architecture. However, in these facultative social-living species, solitary individuals are typically younger individuals that have not yet founded a social group, raising the possibility that the observed enrichment is partly attributable to ontogenetic effects. The social environment is known to modulate neurogenesis in vertebrates: experimentally imposed social isolation has been shown to reduce cell proliferation in both mammals and teleosts, while subordinate teleosts also exhibit reduced neurogenesis (Lieberwirth et al., 2012; Teles et al., 2023; Tunbak et al., 2020). Although naturally occurring solitary fish are unlikely to experience the chronic social stress associated with experimental isolation, differences in social environment are nevertheless expected to shape neural plasticity, as social interactions are potent modulators of gene expression, neurogenesis, and circuit remodeling across vertebrates. The signature of active cell cycle progression and structural dynamics observed in solitary fish may represent a state of proactive neural wiring or sensory priming, optimizing the brain for independent environmental navigation rather than active social management. Together, these findings suggest that social and solitary phenotypes are characterized by alternative modes of neural investment. Rather than representing independent transcriptional programs, the repeated enrichment of biosynthetic and energetic pathways in social-living individuals, alongside structural remodeling and neurodevelopmental pathways in solitary individuals, is consistent with a conserved trade-off in resource allocation within the telencephalon. Maintaining a socially active neural state likely requires sustained investment in protein synthesis, mitochondrial activity, and cellular metabolism to support continuous processing of complex social information. Conversely, when these immediate biosynthetic demands are reduced, greater transcriptional investment may be directed toward structural plasticity, circuit remodeling, and maintenance of neural architecture. Although transcriptomic data cannot directly quantify energetic allocation, the remarkable consistency of this reciprocal pattern across both behavioral plasticity and independent evolutionary transitions suggests that energetic constraints may represent a fundamental mechanism linking neural function to the evolution of social behavior.

The absence of similarly conserved patterns in the diencephalon in relation to social-living highlights that not all components of the Social Decision-Making Network contribute equally to evolutionary convergence. The diencephalon integrates endocrine, metabolic, and motivational signals that regulate behavioral state transitions (Goodson, 2005; Ogawa et al., 2021; Trudeau, 2022), but its heterogeneous organization may obscure conserved transcriptional responses when entire brain regions are analyzed. Alternatively, endocrine regulation of social behavior may have evolved through more lineage-specific molecular mechanisms than those operating in telencephalic circuits responsible for information integration and behavioral decision-making. Future studies combining spatial transcriptomics or single-cell-based approaches with finer anatomical resolution will be necessary to distinguish between these possibilities.

The extensive functional convergence observed between the two independent origins of social living provides strong support for the behavioral genetic toolkit hypothesis. This framework, well established in evo-devo (Carroll, 2008), proposes that repeated evolution of similar behavioral phenotypes occurs through the recurrent co-option of conserved ancestral gene regulatory networks rather than through independent evolution of entirely novel genetic mechanisms (Rittschof & Robinson, 2016; Toth & Robinson, 2007). Although the two facultative species showed virtually no overlap in individual differentially expressed genes, they repeatedly enriched the same biological processes and functional modules, indicating conservation at the level of regulatory architecture rather than individual *loci*. Evidence supporting the existence of a social behavioral genetic toolkit is slowly emerging. Territorial intrusion elicits common functional responses and regulates homologous transcription factors in the brain of mammals, fish and insects (Rittschof et al., 2014). Similar patterns of shared transcriptional programs have been described in the repeated evolution of aggressive behavior among shell-dwelling cichlids (Schumer et al., 2011) and in the convergent evolution of social monogamy across distantly related vertebrates, where independent lineages exhibit highly conserved neural transcriptional signatures despite substantial evolutionary divergence (Young et al., 2019). Our findings therefore extend growing evidence that conserved molecular programs can repeatedly generate similar social phenotypes while allowing species-specific modifications in the genes that regulate those programs.

While our study provides powerful insights into the neurogenomic architecture of sociality, our transcriptomic profiling relied on bulk tissue homogenates from broad brain regions (the whole telencephalon and diencephalon). Because both regions comprise highly heterogeneous sub-nuclei with functionally distinct roles in the Social Decision-Making Network this macro-level approach may obscure cell-type-specific or nuclei-isolated transcriptional responses. This is particularly relevant in the brain, where small neuronal populations can drive major changes in social behavior (Kareklas et al., 2025). Consequently, localized or cell-specific expression changes may have been diluted, partially explaining the low number of individual differentially expressed genes detected despite the robust network-level patterns revealed by GSEA, and the weak association of the diencephalon to the phenotype. Future studies using single-cell approaches combined with spatial transcriptomics will be essential to resolve the cellular and anatomical origins of the conserved transcriptional signatures identified here.

In summary, our study demonstrates that repeated transitions to social living in Lake Tanganyika cichlids are accompanied by the repeated recruitment of conserved transcriptional programs in the telencephalon. Moreover, the same functional modules are deployed during behavioral plasticity within species, suggesting that evolutionary change and behavioral flexibility share a common neurogenomic basis. Rather than relying on a small number of "sociality genes", social evolution appears to emerge through coordinated regulation of ancient gene networks controlling cellular metabolism, protein homeostasis, and neural structural plasticity. These findings provide strong support for the existence of a conserved behavioral genetic toolkit underlying the repeated evolution of sociality while illustrating how polygenic regulatory networks enable both evolutionary innovation and behavioral flexibility.

## Acknowledgements

The authors wish to thank Dr. Magda C. Teles for kindly teaching and assisting during the brain dissections. We would like to thank the Genomics Facility of the Gulbenkian Institute for Molecular Medicine (GIMM) for their sequencing support and technical assistance. We would also like to thank Max Haberbosch, Paul Nuehrenberg, Shun Satoh, and Ryoichi for the kind help collecting underwater individuals and great predisposition during the entire sampling process.

## Data Accessibility

All raw sequencing data have been deposited in the European Nucleotide Archive (ENA; https://www.ebi.ac.uk/ena/browser/home) under the study accession number PRJEB111213. Differential gene expression results are included in Supplementary Data 2 and 3. Custom scripts developed for data processing and analysis are available at https://github.com/psorigue/Group-Living-Evolution, with a permanently archived version accessible via Zenodo (DOI: 10.5281/zenodo.21208439).

## Author Contributions

R.O. and P.S. planned the study; A.J. and P.S. performed the sampling and field work; P.S. performed the bioinformatic analysis and drafted the manuscript. All authors contributed to the discussion of the data and have read and approved the manuscript.

