## Supplementary Figures for "Shared brain transcriptome signatures of social evolution and plasticity in shell-dwelling cichlid fishes"

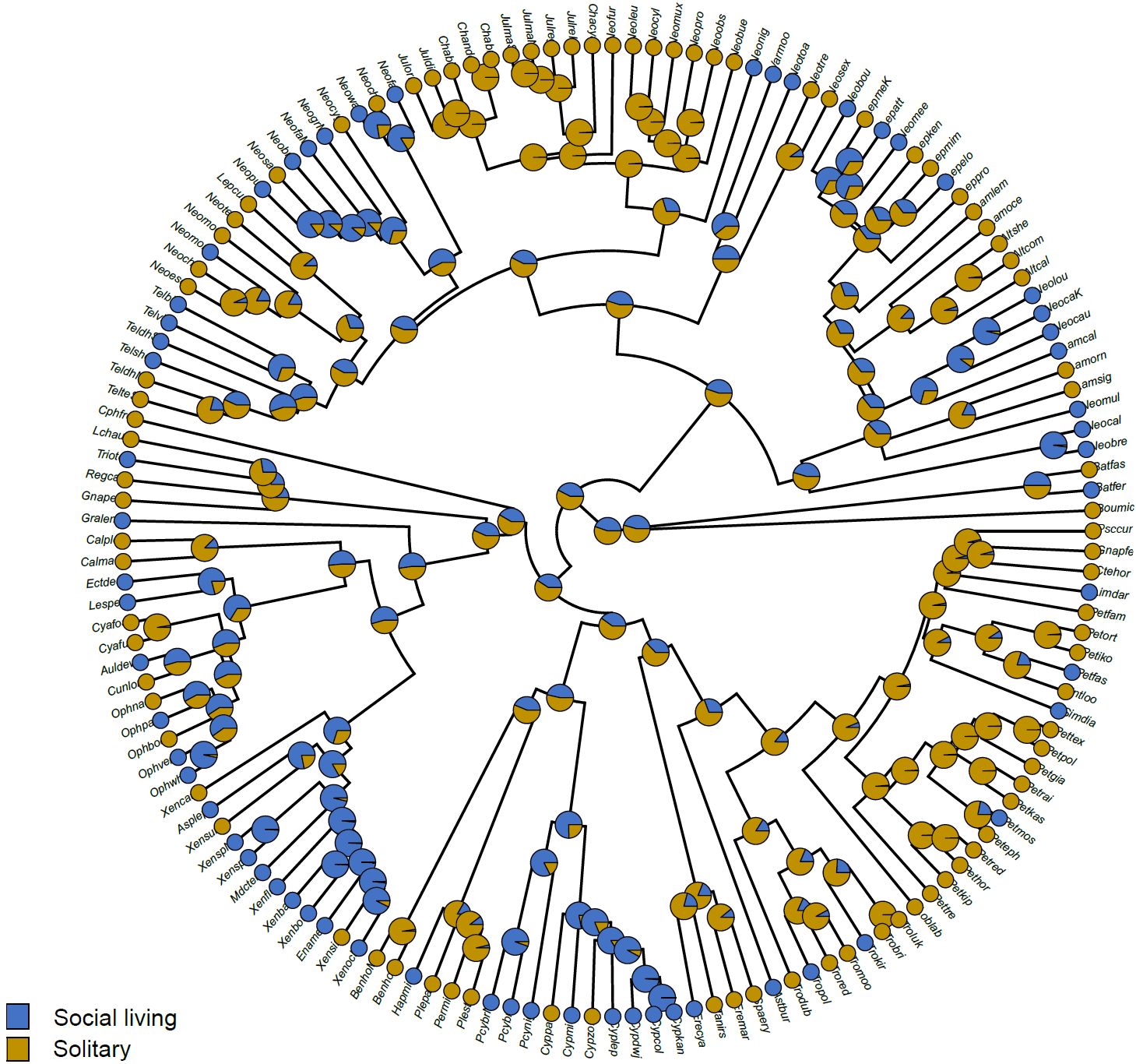

**Supplementary Figure 1: Maximum likelihood ancestral state reconstruction of social organization across the Lake Tanganyika cichlid adaptive radiation.** Ancestral states were reconstructed under an equal-rates (ER) model of discrete character evolution using maximum likelihood implemented in the *ace* function of the *ape* package. The ER model was selected over the all-rates-different (ARD) model based on model comparison (ΔAIC = −0.29; likelihood ratio test, *P* = 0.13). Terminal taxa are classified according to their inferred social organization based on the unsupervised phenotypic characterization of group size. Pie charts at terminal and internal nodes represent the marginal likelihood of each state, with gold indicating solitary and blue indicating social-living. Pie chart proportions correspond to the relative probability of each state, with intermediate proportions reflecting uncertainty in ancestral state estimation. Species aliases were taken from Ronco et a. (2021).

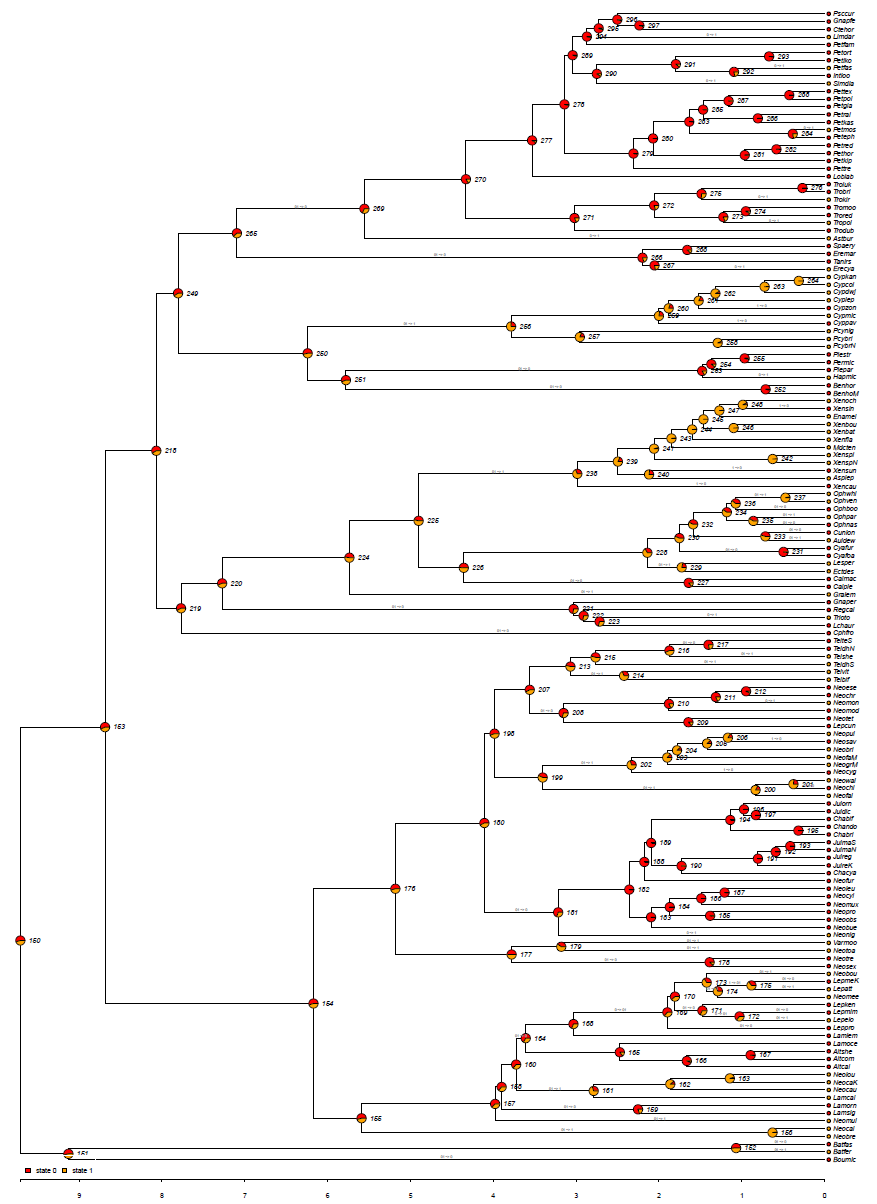

**Supplementary Figure 2: Bayesian ancestral state reconstruction of social organization across the Lake Tanganyika cichlid adaptive radiation.** Ancestral states were reconstructed with the MBASR package in R. Terminal taxa are classified according to their inferred social organization based on the unsupervised phenotypic characterization of group size. Pie charts at terminal and internal nodes represent the marginal likelihood of each state, with red indicating solitary (state 0) and yellow indicating social-living (state 1). Pie chart proportions correspond to the relative probability of each state, with intermediate proportions reflecting uncertainty in ancestral state estimation. Species aliases were taken from Ronco et a. (2021).

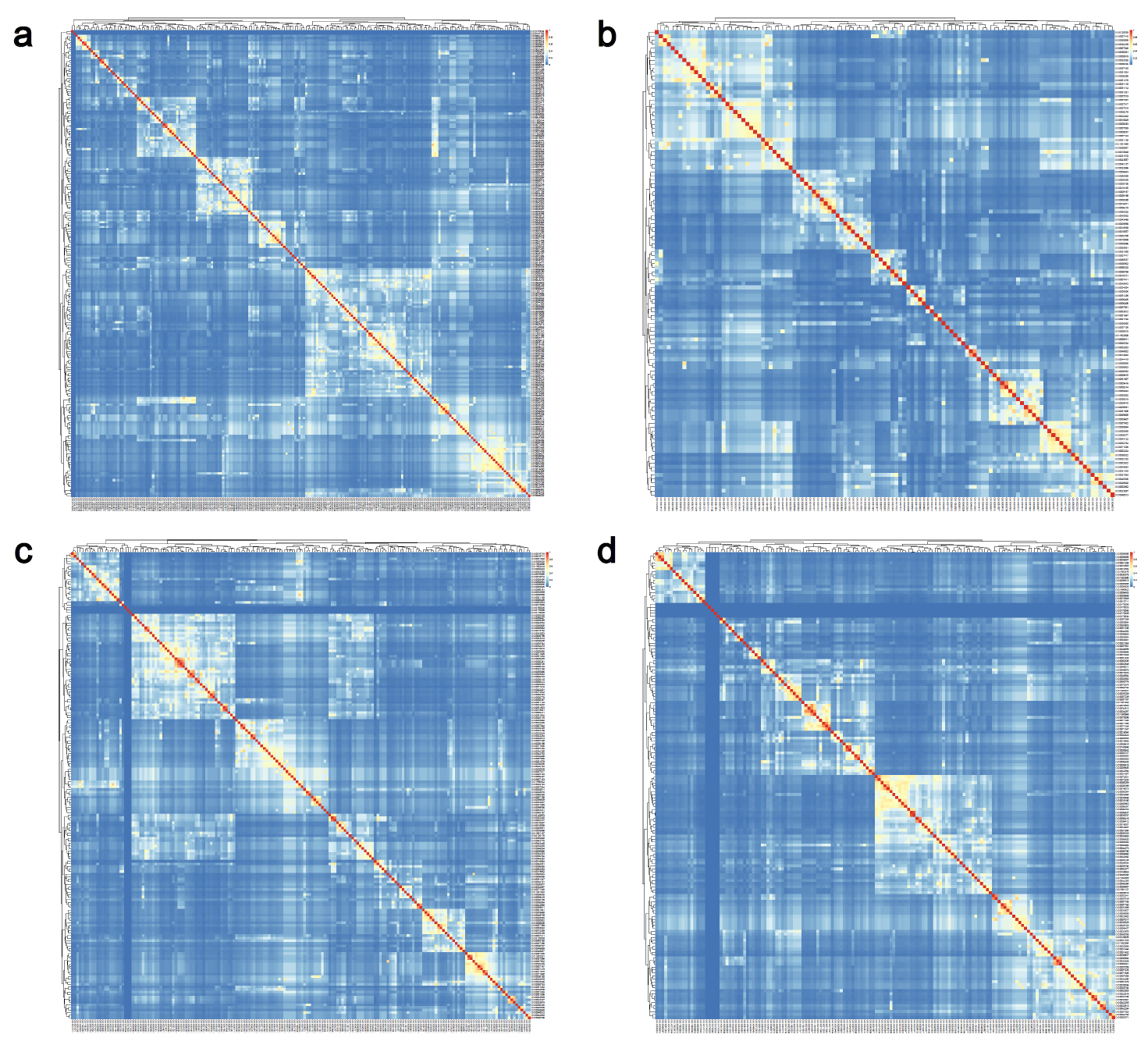

**Supplementary Figure 3. Semantic similarity matrices of enriched Gene Ontology (GO) Biological Process terms identified by Gene Set Enrichment Analysis (GSEA).** Pairwise semantic similarity among significantly enriched GO Biological Process terms was calculated using the Wang semantic similarity measure implemented in the *GOSemSim* R package. The resulting similarity matrices were hierarchically clustered using average linkage with a distance metric of 1 − semantic similarity, and rows and columns were ordered according to the resulting dendrograms. Warmer colors indicate higher semantic similarity between GO terms, whereas cooler colors indicate lower similarity. Clusters were defined by cutting the dendrogram at a height of *h* = 0.75. (a) Telencephalon, intraspecific comparisons; (b) diencephalon, intraspecific comparisons; (c) telencephalon, interspecific comparisons; and (d) diencephalon, interspecific comparisons. The complete lists of GO Biological Process and similarity matrices are provided in the Supplementary Data 3.
